# Observing plasticity of the auditory system: Volumetric decreases along with increased functional connectivity in aspiring professional musicians

**DOI:** 10.1101/2020.11.24.393751

**Authors:** Elisabeth Wenger, Eleftheria Papadaki, André Werner, Simone Kühn, Ulman Lindenberger

## Abstract

Playing music relies on several sensory systems and the motor system, and poses strong demands on control processes, hence, offering an excellent model to study how experience can mold brain structure and function. While most studies on neural correlates of music expertise rely on cross-sectional comparisons, here we compared within-person changes over time in aspiring professionals intensely preparing for an entrance exam at a University of the Arts to skilled amateur musicians not preparing for a music exam. In the group of aspiring professionals, we observed gray-matter volume decrements in left planum polare, posterior insula, and left inferior frontal orbital gyrus over a period of about six months that were absent among the amateur musicians. At the same time, the left planum polare, the largest cluster of structural change, showed increasing functional connectivity with left and right auditory cortex, left precentral gyrus, left supplementary motor cortex, left and right postcentral gyrus, and left cingulate cortex, all regions previously identified to relate to music expertise. In line with the expansion–renormalization pattern of brain plasticity (Wenger, Brozzoli, et al. 2017), the aspiring professionals might have been in the selection and refinement period of plastic change.

---

Playing a musical instrument is an intense, multisensory experience. As music itself is a highly complex stimulus and musicians typically devote a lot of time to their training, they offer an excellent model for studying experience-dependent plastic changes in the brain. In our view, brain plasticity is an adaptive process that is triggered by a prolonged mismatch between the functional supply the brain structure can momentarily provide and the experienced demands the environment poses (Lövdén et al. 2010). In the last years, evidence for malleability of the adult brain structure to environmental challenges has been accumulating, mainly using magnetic resonance imaging (MRI; Lövdén et al. 2013). It has been shown repeatedly that brain volume and number of cells in animals differ depending on their living conditions, for instance, when comparing enriched versus standard rearing environments (Freund et al. 2013). In humans, various challenges like learning how to juggle (Draganski et al. 2004), extensive studying (Draganski et al. 2006), becoming a taxi-driver (Woollett and Maguire 2011), playing a video game (Kühn et al. 2014), or practicing tracing and writing with your non-dominant hand (Wenger, Kühn, et al. 2017) have been shown to elicit changes in estimates of gray-matter volume.

Music expertise has served as a particularly rich and fruitful domain for investigating plastic changes. It involves several sensory systems and the motor system, and it poses high demands on cognitive control processes (Münte et al. 2002; Jäncke 2009; Herholz and Zatorre 2012; Schlaug 2015). It is now well established that musicians typically show an enlargement of brain areas associated with music-related processes in the auditory, motor, and visuospatial domain (Schneider et al. 2002; Gaser and Schlaug 2003; Hutchinson et al. 2003; Bermudez et al. 2009; James et al. 2014). Several brain areas, including the auditory cortices, the anterior corpus callosum, the primary hand motor area and the cerebellum, differ in their structure and size between musicians and control subjects (Münte et al. 2002) and these volumetric differences have been shown to be of behavioral relevance (Schneider et al. 2002; Hyde et al. 2009; Foster and Zatorre 2010a). Groussard and colleagues (Groussard et al. 2014) have identified regions in the brain that increased in volume with the duration of practice, namely left hippocampus, right middle and superior frontal regions, right insula and supplementary motor area, left superior temporal and posterior cingulate areas. Interestingly, while in some regions changes in volume seem to have occurred during early stages of musical training, like in left hippocampus and right middle and superior frontal areas, changes in other areas, specifically in left posterior cingulate cortex, superior temporal areas and right supplementary motor area and insula, were more pronounced or even only occurred after several additional years of practice (Groussard et al. 2014). Similarly, James and colleagues have sorted music expertise into three levels to investigate its influence on gray-matter density (James et al. 2014). While they found gray-matter increases with expertise in areas implicated in working memory and attentional control, that is in fusiform gyrus, mid orbital gyrus, inferior frontal gyrus, intraparietal sulcus, cerebellum, and Heschl’s gyrus, they detected gray-matter decreases with expertise in areas related to sensorimotor function, namely in perirolandic and striatal areas.

Arguably, musicians brains do not only differ structurally from nonmusicians but show also functional differences, such as strengthened functional coupling among relevant regions while performing musical tasks (Herholz and Zatorre 2012). Indeed, numerous functional imaging studies have compared musicians and nonmusicians and have observed differences in activity across many brain regions when individuals were performing musical tasks involving discrimination (Koelsch et al. 2005; Foster and Zatorre 2010b), working memory (Gaab et al. 2006), or production (Bangert et al. 2006; Kleber et al. 2010). Despite the many differences among the tasks used, one area that has been commonly activated in many of these studies was the left superior temporal gyrus, a region that has been linked to musical training in terms of cumulative practice hours (Ellis et al. 2012). Of interest, fMRI studies of perceptual learning with pitch tasks have resulted in both increases (Gaab et al. 2006) and decreases (Jäncke et al. 2001) of activity in auditory areas. Similarly, training to discriminate between melodies constructed of increasingly smaller intervals well below a semitone has been shown to be accompanied by general activation decrements in auditory regions, along with activation increases in frontal cortices (Zatorre, Delhommeau, et al. 2012). Before training, the data had shown the expected dose-response function of more activity with increasing microtonal pitch interval size. After training, however, there was a reduction in blood oxygenation in response to increasing interval size (Zatorre, Delhommeau, et al. 2012), suggesting that learning might decrease the number of neuronal units that are needed to perform the task (Poldrack 2000; Makino et al. 2016).

The brain exhibits spontaneous and systematic activity during wakeful rest (Biswal et al. 1995; van den Heuvel and Hulshoff Pol 2010; Zuo and Xing 2014). Exploiting this characteristic, one can compute resting-state functional connectivity which is based on spontaneous low-frequency fluctuations (< 0.1 Hz) in the blood oxygen level-dependent signal (Biswal et al. 1995), and uncover functional networks that consist of brain regions frequently working together. Activity in the resting state may therefore reflect the repeated history of coactivation within or between brain regions for efficient task performance (e.g., Baldassarre et al., 2016; Cole, Yarkoni, Repovš, Anticevic, & Braver, 2012; Ventura-Campos et al., 2013). Only a few studies have investigated differences in functional connectivity as a function of musical training. Pianists were found to show greater functional connectivity between left auditory cortex and the cerebellum than control participants (Luo et al. 2012). Regions with increases in gray matter in musicians compared to nonmusicians located in posterior and middle cingulate gyrus, left superior temporal gyrus and inferior orbitofrontal gyrus have been shown to have increased connectivity to right prefrontal cortex, left temporal pole, left premotor cortex and supramraginal gyri (Fauvel et al. 2014). Palomar-García and colleagues tested for differences between musicians and nonmusicians in auditory, motor, and audiomotor connectivity and found stronger connectivity between right auditory cortex and right ventral premotor cortex, which correlated with years of practice (Palomar-García et al. 2017). They also found reduced connectivity between motor areas that control both hands in those musicians whose instrument required bimanual coordination, and increased volume in right auditory cortex. This increased gray matter volume correlated negatively with age at which training had begun and was related to increased connectivity between auditory and motor systems (Palomar-García et al. 2017).

As summarized above, most studies on neural correlates of music expertise rely on crosssectional comparisons, rendering conclusions of whether observed group differences were pre-existing or the result of learning de factor impossible. It has been impressively shown, though, that monozygotic twins, i. e. with identical genes, differing on musical training do indeed exhibit neuroanatomical differences, thereby providing strong support for the causal effects of training (de Manzano and Ullén 2018). Still, longitudinal studies with observations within the same individuals over time provide the most direct evidence for effects of musical training on neuroanatomy. We therefore used a variety of methodologies to characterize within-person changes *over time* in aspiring professionals intensely preparing for an entrance exam at a University of the Arts and compared these to skilled amateur musicians not preparing for a music exam. Specifically, we used anatomical MRI along with resting-state fMRI to investigate structural changes in gray-matter volume that arise during this intense learning period within individuals over time and to analyze the changes in functional interactions that accompany these structural changes. We hypothesized that (1) in comparison to amateur musicians, aspiring professional musicians will show volumetric changes in regions previously identified to be relevant in the context of musical training, especially auditory cortex, (2) the regions of structural change will exhibit increased functional connectivity to other regions related to the auditory network, specifically, temporal regions, motor regions, and cingulate gyri and (3) these changes in structure and functional connectivity will be related to behavioral performance.

## Materials and Methods

### Participants

We recruited a total of 24 young adults between 18 and 31 years (M_age_ = 21.92, SD_age_ = 3.72) who were participating in preparatory courses offered at Berlin music schools to prepare them for an entrance exam at a University of Arts. These participants were either aspiring to study to become a conductor, composer, Tonmeister, or instrumentalist. As a control group, we recruited 17 amateur musicians between 18 and 27 years (M_age_ = 23.12, SD_age_ = 3.43) with at least five years of formal music education who were actively performing music in their daily lives but had no aspirations to perform music professionally. All participants had normal hearing, normal or corrected-to-normal vision, no history of psychological or neurological diseases, and no contraindication to participate in an MR study, such as metallic implants, tinnitus, or claustrophobia. The groups did not differ with respect to age (*t*(39) < −1.05, *p* = .30) or years of playing a primary instrument (*t*(38) ^1^ < 1, *p* = .68; aspiring professionals: M_years_ = 12.04, SD_years_ = 4.57; amateur musicians: M_years_ = 12.74, SD_years_ = 5.97).

Participants were paid up to 200€ for completion of the whole study (including up to 5 measurement time points with 1.5 h of MRI and 1.5 h of behavioral testing). The ethical board of the DGPs (Ethikkommission der Deutschen Gesellschaft für Psychologie) approved the study, and written consent of all participants was obtained prior to investigation.

### Experimental Design

Participants were invited for behavioral testing as well as magnetic resonance imaging (MRI) assessment between one to five times, depending on their availability, in the course of about a year, with approximately 10-12 weeks distance between appointments (see Figure 1). Participants were put in the MR scanner for about an hour and 15 min, and were then tested on the in-house developed “Berlin Gehörbildung Scale” (Lin et al. under review), a test to assess music aptitude at expert levels.

**Figure 1.**
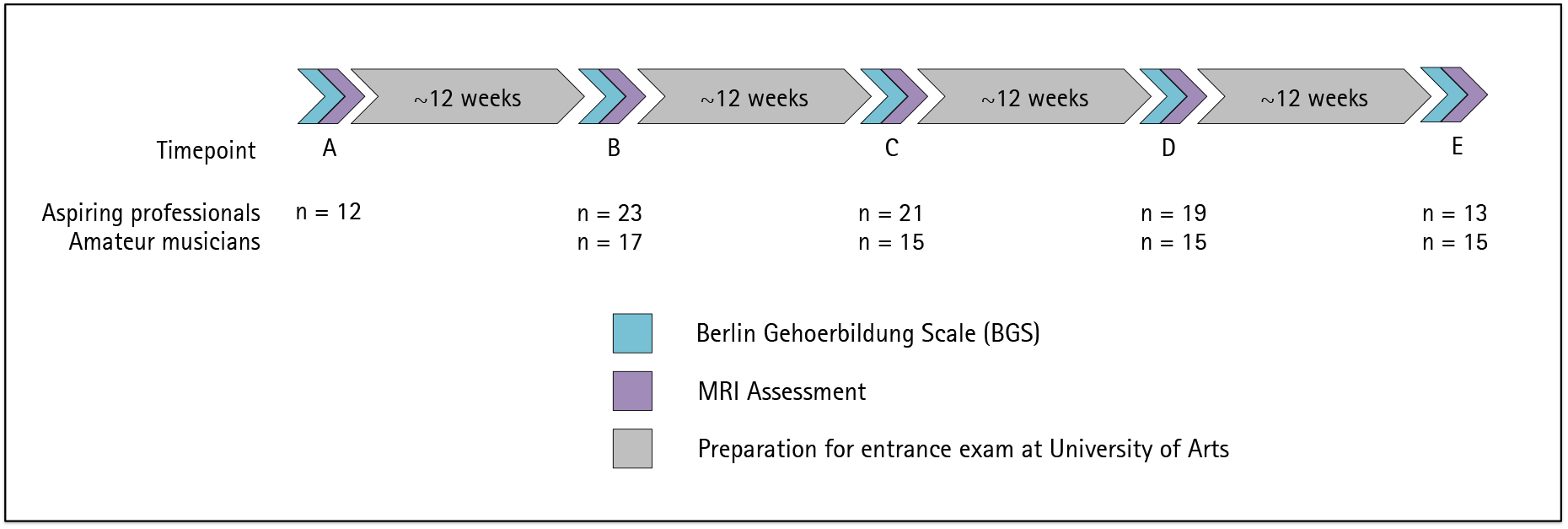
Overview of experimental design with recruitment numbers for aspiring professionals and amateur musicians at each timepoint.

**Figure 2.**
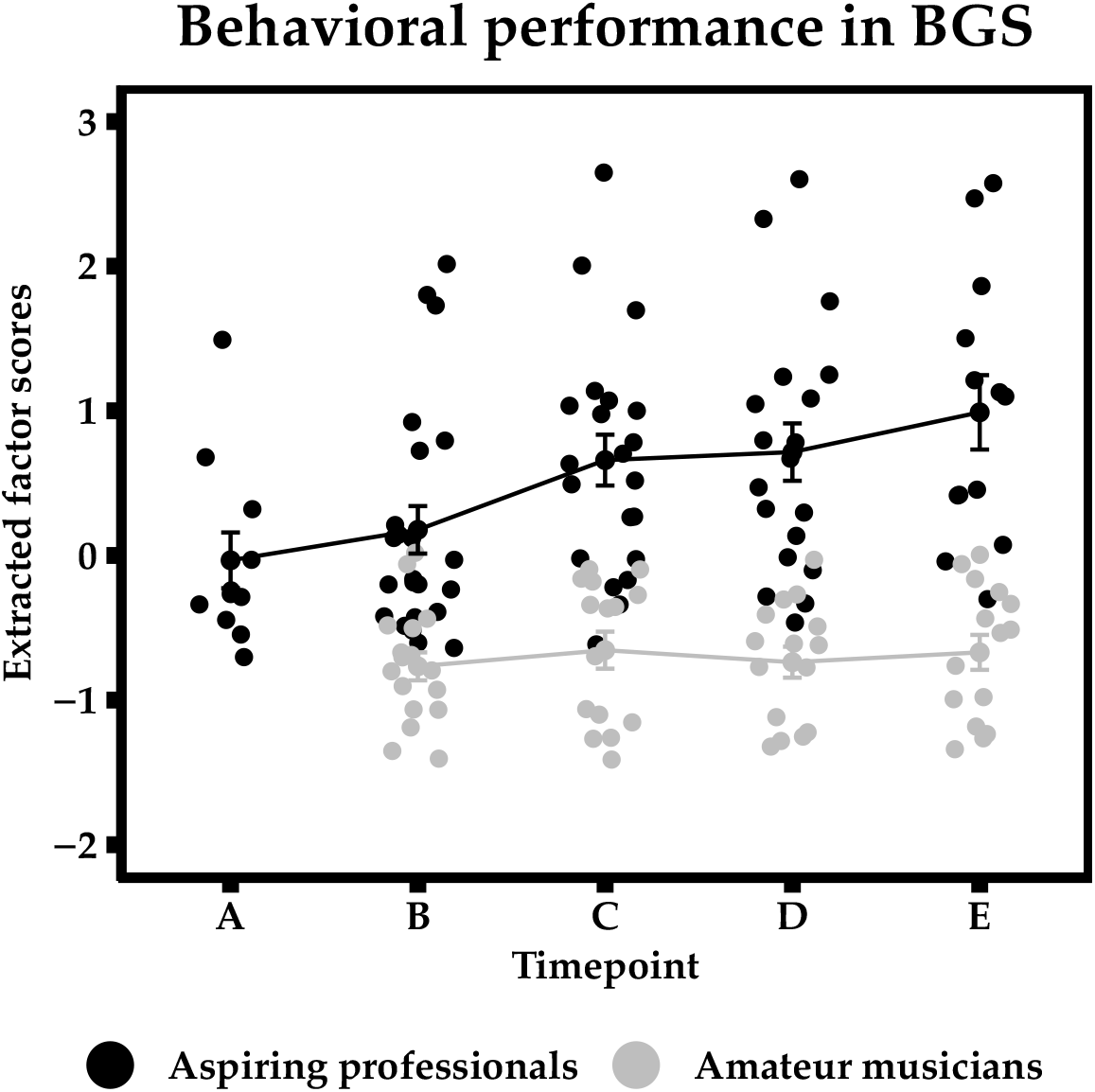
Behavioral performance scores on Berlin Gehoerbildung Scale (BGS). Error bars represent ±1 standard errors (SE).

### Behavioral Measure of Music Expertise

The *Berlin Gehoerbildung Scale* (BGS) was designed by André Werner, a composer and collaborator in this study. It is a listening and transcription task focused on assessing music expertise (for a detailed description see Lin et al. 2020). It is informed by music theory and uses a variety of testing methods in the ear-training tradition. Items cover a variety of topics in music theory and ear training, including intervals, scales, dictation, rhythm, chords, cadences, identifying mistakes in music excerpts, and instrument recognition. Using behavioral data of amateur musicians, aspiring professional musicians, as well as 19 music students already studying music at a University of Arts, we have established a hierarchical structural equation model (SEM) of their behavioral performance the first time they encountered the test (Lin et al. 2020). The hierarchical model postulates four first-order factors of musical abilities, namely “Interval and Scales,” “Dictation,” “Chords and Cadences,” and “Complex Listening,” which together define a second-order factor of general music expertise. These four first-order factors load highly onto the second-order factor music expertise. We fixed the factor loadings of this established model and then extracted the second-order factor scores for each individual at each time point to investigate changes in performance over time. We then entered the factor scores into a repeated-measures ANOVA with the factors Time (timepoint B, C, and D, as these measurement occasions provide us with the largest sample) and Group (aspiring professionals vs. amateurs).

### MRI Data Acquisition

MR images were collected on a Siemens Tim Trio 3T MR scanner (Erlangen, Germany) with a standard 12-channel head coil. The MR measurement protocol included a T1-weighted structural scan and a resting-state acquisition.

As structural images, we used a three-dimensional T1-weighted magnetization prepared gradient-echo sequence (MPRAGE) of 9.20 minutes with the following parameters: TR = 2500 ms, TE = 4.77 ms, TI = 1100 ms, flip angle = 7°, bandwidth = 140 Hz/pixel, acquisition matrix = 256 × 256 x 192, isometric voxel size = 1 mm^3^. We used the prescan normalize option and a 3D distortion correction for non-linear gradients.

Whole brain functional images were collected using a T_2_^*^-weighted EPI sequence of 8 minutes sensitive to BOLD contrast (TR = 2000 ms, TE = 30 ms, FOV = 216 × 216 × 129 mm^3^, flip angle = 80°, slice thickness 3.0 mm, distance factor = 20%, voxel size = 3 mm^3^, 36 axial slices, using GRAPPA acceleration factor 2). Slices were acquired in an interleaved fashion, aligned to genu-splenium of the corpus callosum.

### Structural Data Analysis

The structural MPRAGE images were processed by means of the Computational Anatomy Toolbox (CAT12; v1247; http://dbm.neuro.uni-jena.de/cat/) for SPM12 (v7219; www.fil.ac.uk/spm/) in Matlab 2017a (the Mathworks, Inc., Natick, MA, USA). Using default parameters, pre-processing of the data involved intra-subject realignment, bias-field and noise removal, skull stripping, segmentation into gray (GM) and white matter (WM) and cerebrospinal fluid (CSF), and finally normalization to MNI space using DARTEL to a 1.5 mm isotropic adult template provided by the CAT12 toolbox (whereby normalization is estimated for the mean image of all time points and then applied to all images). The resulting gray matter (GM) maps were smoothed with a standard gaussian kernel of 8 mm full-width at half maximum (FWHM).

As for quality assurance, images were first visually inspected for artefacts prior to processing. Then, a statistical quality control based on inter-subject homogeneity after segmentation was conducted using the “check homogeneity” function in CAT12. After preprocessing, all images were visually checked again for artifacts, whereby none were detected.

Statistical analysis of the GM maps was first carried out by means of a 2-sample *t*-test to test for initial structural differences between aspiring professional and amateur musicians at measurement occasion B (the first timepoint where both groups were fully recruited). This analysis included 23 aspiring professionals and 17 amateur musicians. An absolute graymatter probability threshold of 0.2 was applied. To control for type-I error, a significant effect was reported when the results met a peak-level threshold of *p* < 0.005 and when the cluster size exceeded the expected voxels per cluster threshold (k > 259 in this case) in combination with correction for non-isotropic smoothness. The expected voxels per cluster threshold was computed automatically by the CAT12 toolbox according to random field theory and empirically determines the minimum number of voxels that, in combination with a voxel-level threshold, clusters must meet in order to be reported (Hayasaka and Nichols 2004). In addition, correction for non-isotropic smoothness adjusts the minimum cluster size depending on the local smoothness of the data. This is a common cluster correction method used for whole-brain VBM analyses.

To further characterize pre-existing structural differences in gray-matter volume between those two groups of musicians, we additionally performed a region-of-interest (ROI) analysis, focusing on left and right superior temporal gyrus, as well as further divisions into bilateral planum temporale, Heschl’s gyrus, and planum polare (taken from the HarvardOxford atlas https://identifiers.org/neurovault.collection:262) (Desikan et al. 2006).

The main analysis in this paper focused on differential changes over time in the two groups of musicians by means of a whole brain flexible factorial design with a focus on the interaction Time x Group. Since not all participants provided data for all time points, we based our statistical analysis on the middle three measurement occasions (B, C, and D) and only included those participants that contributed data to those three time points since this provided us with the highest possible number of participants for a longitudinal analysis in SPM. This resulted in a final sample of 19 aspiring professionals and 15 amateur musicians in this statistical comparison in which we tested for brain regions that display a significant increase or decrease in aspiring professionals compared to amateur musicians over time. Again, an absolute gray-matter probability threshold of 0.2 was applied. To control for type-I error, here, a significant effect was reported when the results met a peak-level threshold of *p* < 0.001 and when the cluster size exceeded the determined expected voxels per cluster threshold (k > 47) in combination with correction for non-isotropic smoothness (as explained above).

To investigate potential relationships between brain volume changes in the clusters showing a significant Time x Group interaction with behavioral performance, we extracted the data from significant clusters using the REX toolbox (region-of-interest extraction tool; The Gabrieli Lab, MIT; http://www.alfnie.com/software), subtracted pretest from posttest values and correlated the difference scores with behavioral performance scores using Pearson’s correlation coefficient.

### Functional Data Analysis

Data pre-processing of the resting state data was performed using the toolbox DPABI (v4.0) (Yan et al. 2016) running under Matlab2014b. The first 10 EPI volumes were discarded to allow the magnetization to approach a dynamic equilibrium. All volume slices were corrected for different acquisition times and then realigned. Individual structural images were coregistered to the mean functional image after realignment. The transformed structural images were then segmented into GM, white matter (WM), and cerebrospinal fluid (CSF) (Ashburner and Friston 2005). To remove head motion, respiratory and cardiac effects, we regressed out the Friston 24-parameter model (Friston et al. 1996) as well as signals from WM and CSF. In addition, linear and quadratic trends were also included as regressors since the BOLD signal exhibits low-frequency drifts. The DARTEL tool (Ashburner 2007) was used to normalize the functional data to the Montreal Neurological Institute (MNI) template. We used a spatial filter of 4 mm FWHM and finally performed temporal filtering (0.01–0.1 Hz).

We then conducted an exploratory analysis by means of DPABI computing *functional connectivity maps with a seed region* consisting of left planum polare in MNI space, taken from the Harvard Oxford atlas (Desikan et al. 2006). To do so, the mean time course of all voxels in the seed region was used to calculate pairwise linear correlations (Pearson’s correlation) with other voxels in the brain. Individuals’ *r* values were normalized to *z* values using Fisher’s *z* transformation.

Statistical analysis of the functional connectivity maps was again carried out by means of a whole brain flexible factorial design, again focusing on measurement occasions B, C, and D. We entered the images containing the *z*-transformed correlation values (between the seed region planum polare and all other voxels in the brain) in the second-level analysis with a focus on a time-by-group interaction, using a family-wise error (FWE) correction for multiple comparisons at *p* < .05 (cluster size k = 20 voxels). We used the REX toolbox to extract the *z*-transformed correlation coefficient values from within those clusters showing a significant time-by-group interaction.

### Graph Theory Analysis

To perform connectivity analysis using graph-theory measures, we used BRAPH (BRain analysis using GraPH theory) (Mijalkov et al. 2017), a toolbox written in Matlab that uses the Brain Connectivity Toolbox codebase (https://sites.google.com/site/bctnet/) (Rubinov and Sporns 2010) to calculate network matrices. Such correlation matrices based on *r* correlation values were generated for every subject and then utilized in the calculation of both global and nodal measures. In this framework, nodes are brain regions based on the parcellation of the HarvardOxford atlas (Desikan et al. 2006) and edges represent the correlations between the temporal activation of pairs of brain regions. The constructed matrix is a weighted undirected matrix, where the edges indicate the strength of the connection. As is common practice, only positive values were used in the calculation of nodal and global metrics (negative correlations were set to zero).

We computed five *nodal measures* including degree, path length, global efficiency, local efficiency, and the clustering coefficient. The **degree** refers to the total number of edges connected to a node. In the calculations, the weights of the connections were ignored by binarizing the connectivity matrix so that only edges with nonzero weights were considered connected. **Path length** refers to the average distance from a node to all others. The distance between two nodes is defined as the length of the shortest path between those nodes. In the case of a weighted undirected graph, the length of an edge is a function of its weight. Typically, the edge length is inversely proportional to the edge weight (, i.e., a high weight implies a shorter connection). The **global efficiency** at the nodal level defines the efficiency of the information transfer from one region to the whole network, which assesses the average inverse shortest path length between one node and all other nodes in the network. The **local efficiency** as a nodal measure is calculated as the global efficiency of the node on the subgraph level, created by the node’s neighbors. It reflects the efficiency of the information transfer from each region to the neighboring regions. The **clustering** coefficient at a nodal level is calculated as the fraction of triangles present around a node and is a measure of segregation. It reflects the ability for specialized processing in small groups of nodes and is thus regarded a measure of local connectedness within a network.

In addition, we computed four *global measures*, namely characteristic path length, global efficiency, local efficiency and clustering coefficient. The **characteristic path length** as a global measure is calculated as the average of the path lengths of all nodes. **Global efficiency** at the global level is the average of the global efficiency of all nodes in the graph and is inversely related to the characteristic path length. **Local efficiency** computed on the global level is the average of the local efficiencies of its nodes and reflects how well the nodes communicate with adjacent nodes. The **clustering coefficient** as a global metric is the average of the clustering coefficients of all nodes.

Statistical significance testing was done by extracting the values of the three measurement occasions for local and global measures for each subject from BRAPH and then testing for a time-by-group interaction separately for each nodal and global measure using SPSS, in the end applying a correction for multiple comparisons using the false discovery rate (FDR) algorithm (*p*-value of .05; https://www.sdmproject.com/utilities/?show=FDR).

## Results

### Behavioral Results

Based on BGS results, aspiring professional musicians showed significantly higher levels of general music expertise than amateur musicians at measurement occasion B, which corresponds to an early phase of assessment, *t*(32) = 4.57, *p* < .001, Hedges’ *g* = 1.58. Furthermore, aspiring professionals showed an increase in performance, whereas amateurs’ performance remained relatively stable, as reflected by a significant time-by-group interaction, *F*(2,64) = 8.53, *p* = .001, partial η squared = 0.21.

### Preexisting Differences in GM Volume Between Aspiring Professionals and Amateur Musicians

To characterize differences in gray-matter volume between aspiring professionals and amateur musicians, we first computed a 2-sample *t*-test on the segmented whole-brain graymatter maps at measurement occasion B. This cross-sectional comparison yielded four significant clusters in superior parietal lobule, left superior temporal gyrus, right hippocampus, and right postcentral gyrus (see Table 1 and Figure 3), in which participants of the aspiring professional group showed greater gray-matter volume than amateur musicians.

**Table 1.** Brain regions showing a significant group difference in gray-matter volume between aspiring professionals and amateur musicians at measurement occasion B (*p* < .005, nonstationary smoothness corrected and cluster correction for expected voxels).

| Area | Peak coordinates (MNI) | T-score | Extent |
| --- | --- | --- | --- |
| Right Hippocampus | 22 -18 -24 | 3.54 | 567 |
| Right superior parietal lobule | 42 -38 52 | 4.00 | 415 |
| Left superior/middle temporal gyrus | -52 -26 -9 | 3.63 | 348 |
| Right postcentral gyrus | 9 -34 74 | 3.54 | 111 |

**Figure 3.**
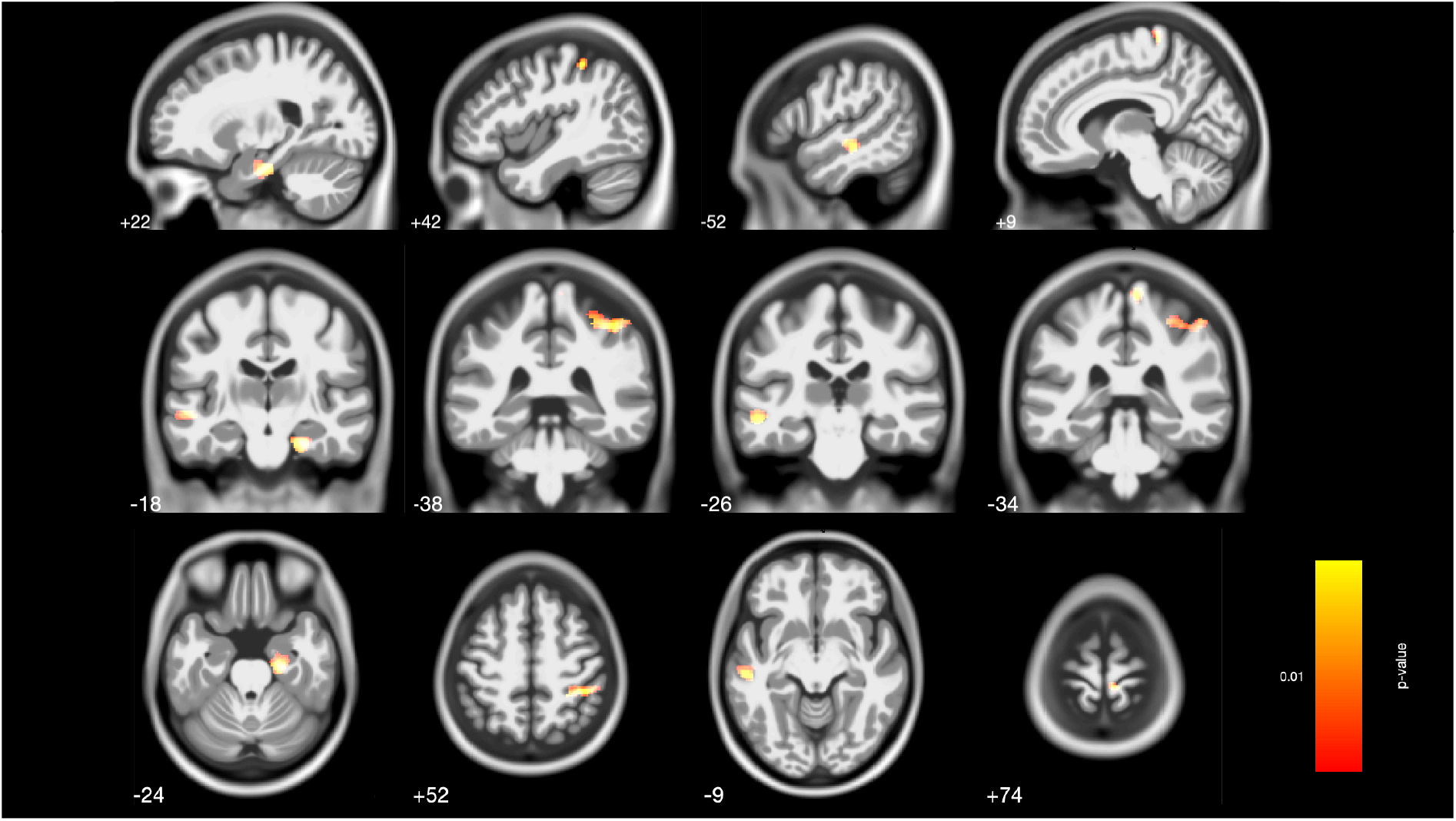
Regions of preexisting differences in gray-matter volume between aspiring professionals and amateur musicians at measurement occasion B in hippocampus, superior parietal lobule, superior/middle temporal gyrus, and postcentral gyrus emerging in a wholebrain 2-sample *t*-test (*p* < 0.005, nonstationary smoothness corrected and cluster correction for expected voxels). Coordinates refer to MNI space. In all cases, volumes were greater in aspiring professionals than in amateur musicians.

An additional ROI analysis, focusing on primary and secondary auditory cortex further confirmed a significant difference in the right anterior portion of superior temporal gyrus (STG) (*t*(38) = 2.40, *p* = .02, *Hedges*’ *g* = 0.7531) and the left posterior portion of STG (*t*(38) = 2.37, *p* = .02, *Hedges*’ *g* = 0.7419) (see Figure 4). Analyses of gray-matter volume differences in bilateral planum temporale, Heschl’s gyrus, and planum polare showed the same tendency of greater gray-matter volumes in aspiring professionals than in amateur musicians but failed to reach the threshold of statistical significance.

**Figure 4.**
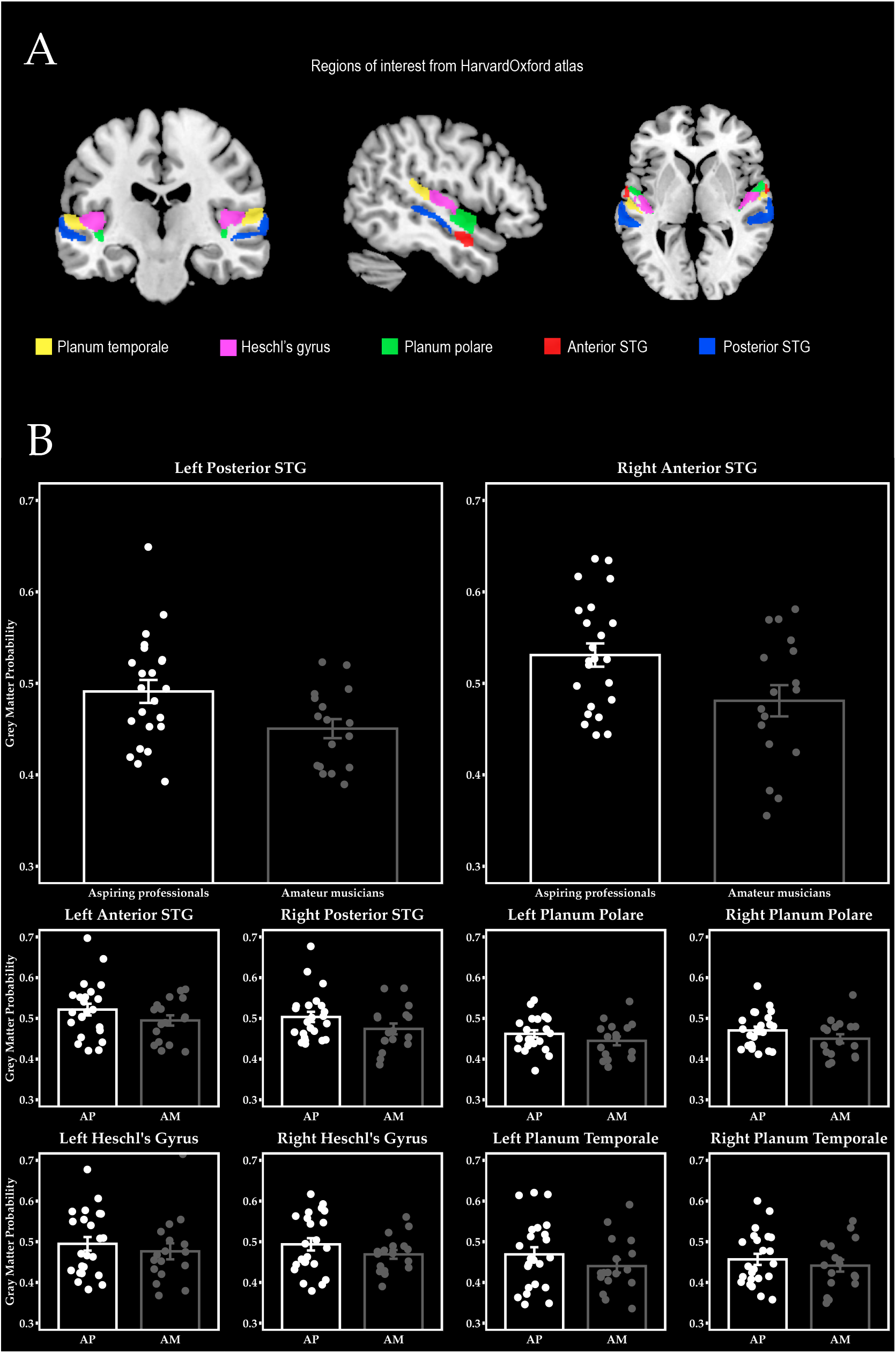
Region-of-interest (ROI) analyses showed a significant difference in gray-matter volume in left posterior superior temporal gyrus (STG) and right anterior STG (*p* < .05). All other ROIs showed the same tendency of greater gray-matter volumes in aspiring professionals than in amateur musicians but failed to reach the threshold of statistical significance.

### Changes in GM Volume over Time

Given that the focus of this study was on differences in within-person changes between aspiring professionals and amateurs, we computed a whole-brain interaction on the segmented whole-brain gray-matter maps. We found three significant clusters, namely in left planum polare, left posterior insula extending into planum polare, and left inferior frontal orbital gyrus extending into anterior insula (see Figure 5 and Table 2 for exact coordinates and *F*-scores). All of these clusters were driven by decreases in gray-matter volume in aspiring professional musicians relative to amateur musicians (see Figure 5b).

**Table 2.** Brain regions showing a significant interaction effect of Group (aspiring professionals vs. amateur musicians) and Time (timepoint B, C and D) in gray-matter volume (*p* < .001, nonstationary smoothness corrected and cluster correction for expected voxels)

| *Area* | *Peak coordinates (MNI)* | *F-score* | *Extent* |
| --- | --- | --- | --- |
| Left planum polare | -48 -14 0 | 23.02 | 292 |
| Left posterior insula / planum polare | -38 -3 -21 | 18.40 | 181 |
| Left inferior frontal orbital gyrus /<br>anterior insula | -32 24 -6 | 16.06 | 181 |

**Figure 5.**
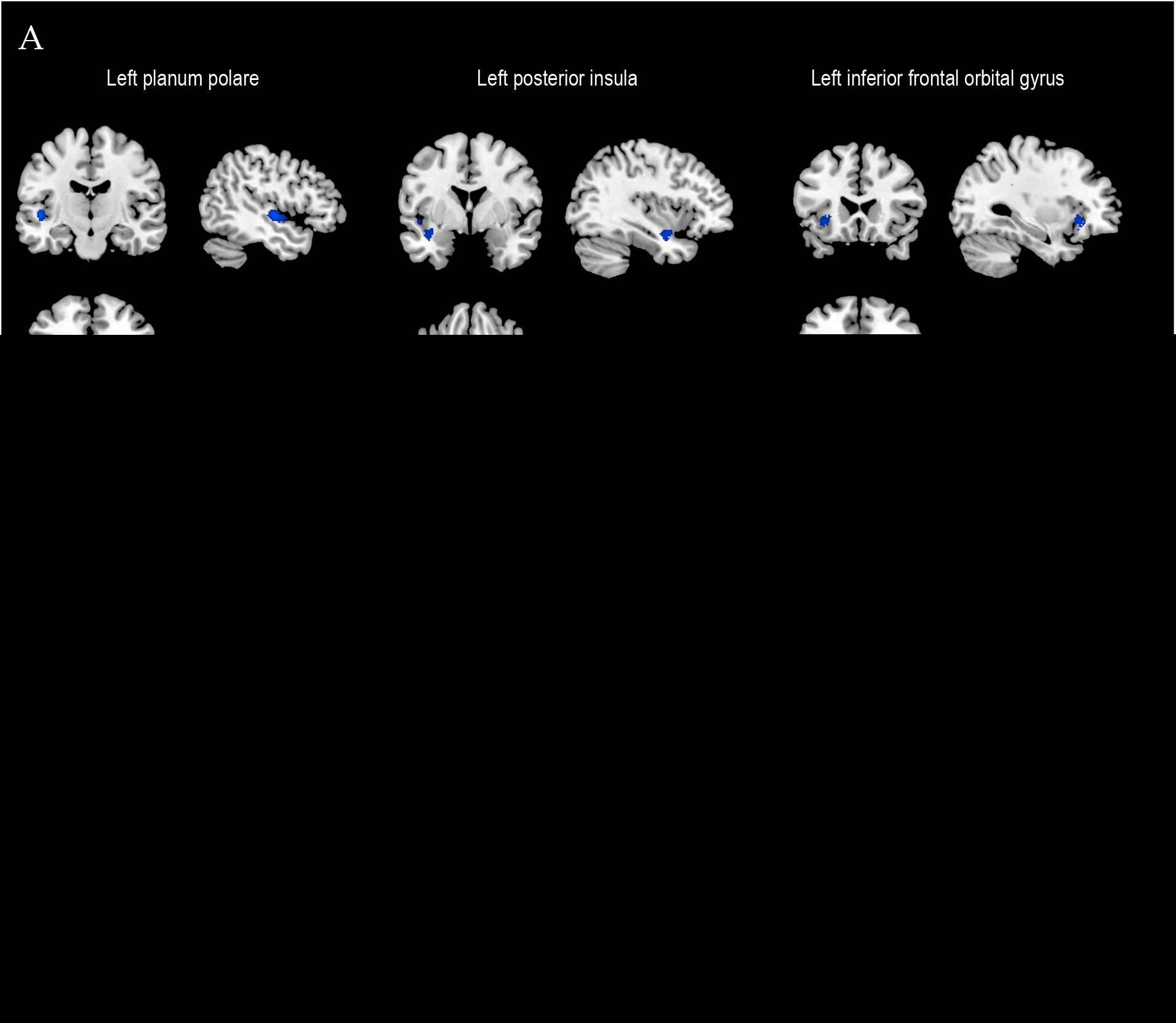
(A) Significant clusters in left planum polare, posterior insula and inferior frontal orbital gyrus emerging in a whole-brain time-by-group interaction analysis (*p* < 0.001, k > 47, corrected for nonstationary smoothness). Coordinates refer to MNI space. (B) Bargraphs with the extracted gray-matter volume estimates of the significant clusters in the time-by-group interaction. This effect is driven by a decrease of gray-matter volume in aspiring professionals compared to amateur musicians. Error bars represent ±1 SE.

For the left planum polare and inferior frontal orbital gyrus (IFoG), the observed decrements in estimates of gray-matter volume in the group of aspiring professionals correlated with general music expertise as assessed by the BGS at measurement occasions B, C, and D (see Figure 6). A similar result was obtained at trend level for the posterior insula (left planum polare: *r*_Time B_ (19) = −0.581*, *p* = .009; *r*_Time C_ (19) = −0.517*, *p* = .023; *r*_Time D_(19) = −0.588*, *p* = .008; left posterior insula: *r*_Time B_ (19) = −0.387, *p* = .102; *r*_Time C_ (19) = −0.525*, *p* = .021; *r*_Time D_ (19) = −0.433, *p* = .064; left IFoG: *r*_Time B_ (19) = −0.558*, *p* = .013; *r*_Time C_ (19) = −0.634*, *p* = .004; *r*_Time D_ (19) = −0.589*, *p* = .008). This association was also true across the whole sample (left planum polare: *r*_Time B_(34) = −0.580*, *p* < .001; *r*_Time C_ (34) = −0.523*, *p* = .001; *r*_Time D_ (34) = −0.599*, *p* < .001; left posterior insula: *r*_Time B_ (34) = −0.282, *p* = .106; *r*_Time C_ (34) = −0.398*, *p* = .020; *r*_Time D_ (34) = −0.373*, *p* = .030; left IFoG: *r*_Time B_ (34) = −0.586*, *p* < .001; *r*_Time C_ (34) = −0.620*, *p* < .001; *r*_Time D_(34) = −0.620*, *p* < .001). Importantly, no such associations were found within the group of amateur musicians (all *p*s > .08).

**Figure 6.**
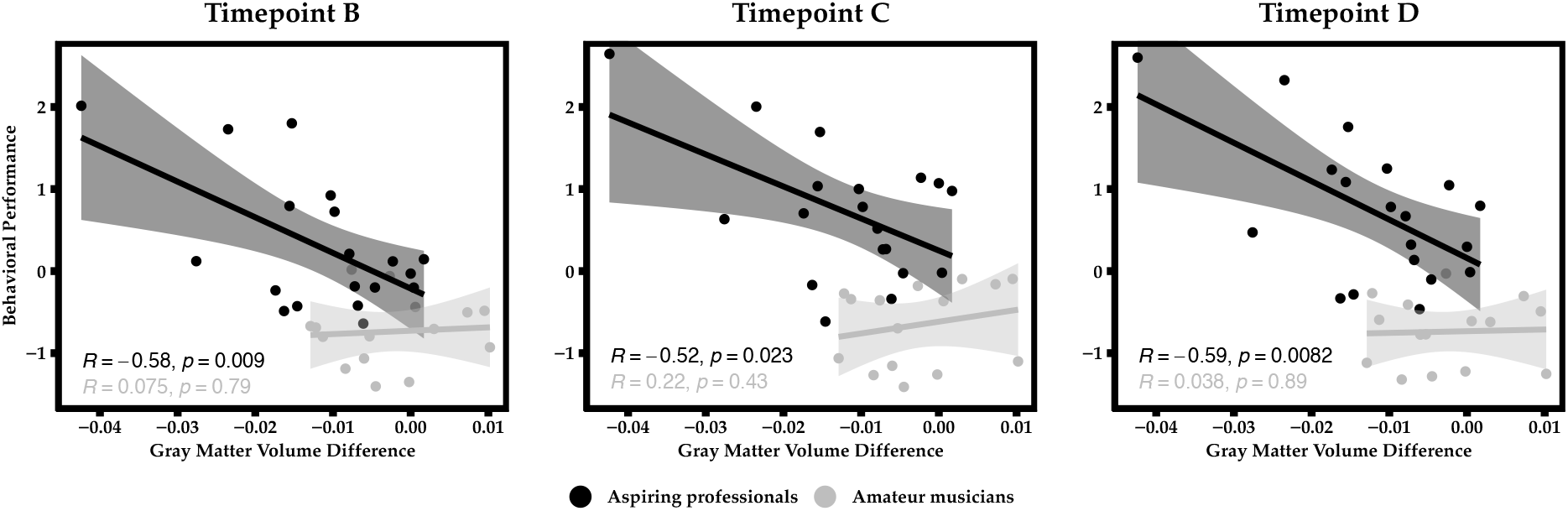
Correlations between decrease in gray-matter volume in left planum polare and behavioral performance in the BGS at measurement occasions B, C, and D.

Correlations in the total sample continued to differ reliably from zero in planum polare and inferior frontal gyrus after excluding one very high-performing individual who also exhibited the most pronounced structural decrease (but does not qualify as an outlier; left planum polare: *r*_Time B_ (33) = −0.434*, *p* = .012; *r*_Time C_ (33) = −0.354*, *p* =.043; *r*_Time D_ (33) = −0.468*, *p* = .006; left IFoG: *r*_Time B_ (33) = −0.519*, *p* = .002; *r*_Time C_ (33) = −0.559*, *p* = .001; *r*_Time D_ (33) = −0.561*, *p* = .001, but not in left posterior insula: *r*_Time B_ (33) = 0.00, *p* = .999; *r*_Time C_ (33) = −0.162, *p* = .367; *r*_Time D_ (33) = −0.141, *p* = .433). This means that those individuals showing the highest proficiency in this behavioral test were also the ones that exhibited the most pronounced decrease in gray-matter volume. In contrast, the decrease in estimates of gray-matter volumes did not correlate with improvements in music expertise (*r*_planum polare_ (19) = −0.104, *p* = .671; *r*_posterior insula_ (19) −161, *p* = .483; *r*_IFoG_ (19) −0.171, *p* = .483).

### Changes in Functional Connectivity

To understand these changes in gray-matter volume, we further investigated training-dependent changes in the coupling between brain regions. Here, we focused on the largest cluster of structural change located in left planum polare, that is, auditory cortex, and its correlations with other regions of the brain. We found increasing functional connectivity of the left planum polare to left and right auditory cortex, left precentral gyrus and left supplementary motor cortex, left posterior cingulate, and left and right postcentral gyrus over time in aspiring professionals compared to amateur musicians (FWE-corrected *p*-value of 0.05; see Figure 7).

**Figure 7.**
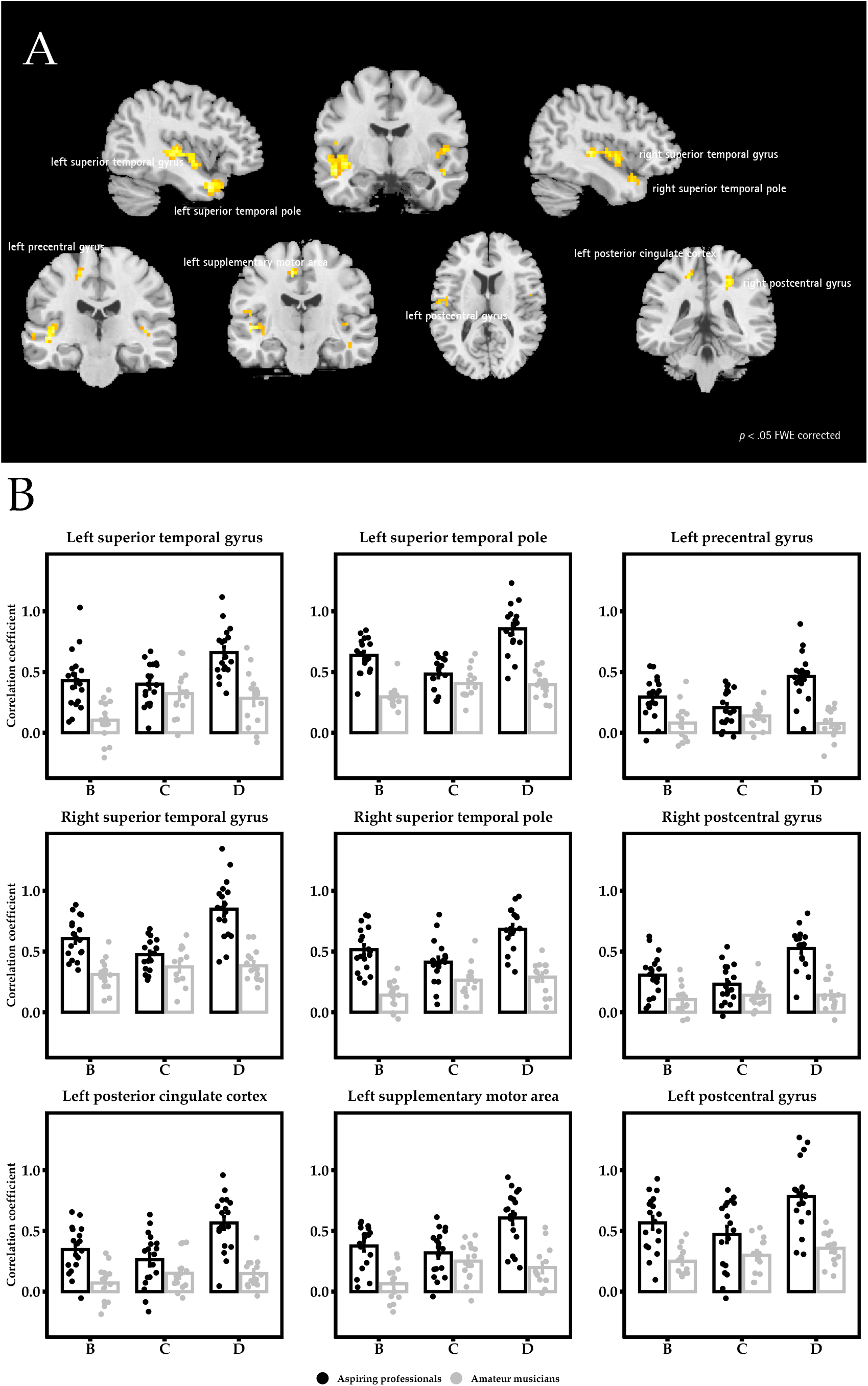
(A) Significant clusters exhibiting increased functional connectivity over time with left planum polare in aspiring professionals compared to amateur musicians (*p* < .05 FWE corrected). (B) Bargraphs with the extracted Fisher’s z-transformed correlation coefficients from those significant clusters of the Time-by-Group interaction. Group-by-time interactions of the functional connectivity analysis were driven by increasing correlation coefficients in aspiring professionals relative to stable correlations among amateur musicians. Error bars represent ±1 SE.

### Changes in Graph-theoretical Measures

To further characterize the network in which left planum polare participates, we conducted graph-theory analyses and compared network characteristics in the two groups over time. While there were no significant time-by-group interactions in any of the nodal measures, there were significant time-by-group interactions for all global measures, namely for characteristic path length, global efficiency, local efficiency, and clustering (see Table 3 for exact numbers). In all of those measures, the group of amateur musicians showed no reliable mean change over time, whereas the group of aspiring professionals showed significant increases over time in all global metrics except for path length, which, as expected, decreased over time.

**Table 3.** Nodal and global measures of graph theoretical analyses at measurement occasions B, C and D, comparing aspiring professionals to amateur musicians.

| Nodal Measures | Aspiring professionals |  |  | Amateur musicians |  |  | *F* | *p* (FDR-corr.) |
| --- | --- | --- | --- | --- | --- | --- | --- | --- |
|  | Time B | Time C | Time D | Time B | Time C | Time D |  |  |
| Degree | 98.83 | 101.16 | 107.33 | 99.85 | 105.21 | 104.78 | 1.61 | .20 |
| Path length | 2.76 | 2.63 | 2.21 | 3.10 | 2.48 | 2.53 | 2.74 | .15 |
| Global efficiency | 0.42 | 0.45 | 0.52 | 0.37 | 0.46 | 0.45 | 2.16 | .15 |
| Local efficiency | 1.97 | 2.24 | 2.89 | 1.54 | 2.28 | 2.13 | 2.95 | .15 |
| Clustering | 0.35 | 0.39 | 0.47 | 0.29 | 0.39 | 0.37 | 2.49 | .15 |

| Global Measures | Aspiring professionals |  |  | Amateur musicians |  |  | *F* | *p* (FDR-corr.) |
| --- | --- | --- | --- | --- | --- | --- | --- | --- |
|  | Time B | Time C | Time D | Time B | Time C | Time D |  |  |
| Characteristic path length | 2.89 | 2.78 | 2.36 | 3.15 | 2.69 | 2.77 | 3.14 | .05 |
| Global efficiency | 0.40 | 0.42 | 0.49 | 0.36 | 0.43 | 0.41 | 3.73 | .05 |
| Local efficiency | 1.84 | 2.03 | 2.65 | 1.47 | 2.06 | 1.92 | 3.94 | .05 |
| Clustering | 0.33 | 0.36 | 0.44 | 0.28 | 0.36 | 0.35 | 3.44 | .05 |

## Discussion

In the present longitudinal study, we set out to investigate structural brain alterations and changes in functional connectivity in musicians intensely preparing for their entrance exam at a University of Arts. We found that gray-matter volume decreased over time in comparison to amateur musicians in three clusters, namely left planum polare, posterior insula extending into planum polare, and left inferior frontal orbital gyrus extending into anterior insula. The biggest cluster of structural change was observed in left planum polare, which exhibited increased functional connectivity with left and right auditory cortex, left precentral gyrus, left supplementary motor cortex, left posterior cingulate cortex, and left and right postcentral gyrus. All of these regions have been previously identified to play important roles in music expertise (e.g., Luo et al. 2012; Groussard et al. 2014). The increase in connectivity for the region showing the greatest structural change was also reflected in results based on graph theory. Here, we observed changes over time in the global metrics, indicating participation of the planum polare in an increasingly complex network in the group of aspiring professionals compared to amateur musicians.

Our results once again speak to the malleability of adult brain structure to environmental influences (Lövdén et al. 2013; Kühn and Lindenberger 2016; Lindenberger et al. 2017). The *left planum polare* as a region within the superior temporal gyrus, adjacent to left Heschl’s gyrus, has been reported to show preferential activity to musical stimuli in comparison to other types of complex sounds, such as speech and non-linguistic vocalizations, and to integrate acoustic characteristics in the context of complex musical sounds, both in trained musicians and non-musicians (Angulo-Perkins et al. 2014). In another study, left planum polare showed activity during high-level musical processing (Brown et al. 2004). In a study looking into functional networks underlying music processing and processing of vocalizations with a passive listening stimulation paradigm that included different vocal sound categories (i.e., song, hum and speech), left planum polare together with planum temporale and a group of regions on the right hemisphere that included the supplementary motor area, premotor cortex and the inferior frontal gyrus, showed stronger activations during music listening (Angulo-Perkins and Concha 2019). Interestingly, left planum polare also showed activity during vocal musical listening, with and without lyrics, a finding pointing towards its role in music processing of temporally complex sounds, such as vocal music and speech. Overall, evidence suggests that the planum polare might be playing an intermediate role between the primary auditory cortex and other associative cortices, possibly extracting information (such as melodic patterns or pitch-interval ratios) required for further processing leading to perceptual evaluations (e.g., a same-different task), vocal production, and sensory-motor coordination to reproduce melodic or rhythmic sounds (Angulo-Perkins and Concha 2019).

As an integration hub, the *insula* serves a plethora of different tasks, including sensory, emotional, motivational and cognitive functions (Gogolla 2017). More specifically within the realm of music, the insula has often been discussed to reflect the emotional aspects of music processing (Blood and Zatorre 2001; Koelsch et al. 2005; Koelsch 2010) and is involved in autonomic regulation and sensory representation of emotion percepts (Koelsch 2014). As aspiring professional musicians do not only have to perfect their technical skills but also have to hone their emotional sensitivity to music, it is conceivable that insula cortex, both anterior and posterior portions, evinces structural change.

*Left inferior frontal gyrus* is well known for its role in syntactic processing of language and music (Friederici 2002; Tillmann et al. 2006; Nan and Friederici 2012), as well as more broadly in general cognitive functions, such as top-down attention and working memory (Janata et al. 2002; Schulze et al. 2011). Especially the orbitofrontal part has been associated with automatic appraisal and is activated by breaches of expectancy (Koelsch 2014), a function crucial for aspiring professional musicians, as it helps them to discriminate, for instance, between expected and unexpected chord progressions. Interestingly, there have been findings of projections from the anterior superior temporal plane to the orbitofrontal cortex in rhesus monkeys (Petrides and Pandya 1988), that go along well with a recent finding of functional connectivity of the left planum polare with orbitofrontal cortex in an fMRI study during music-evoked emotional processing (Koelsch et al. 2018).

Within all three of these regions, we have found structural decreases in the group of aspiring professionals, while volumes in amateur musicians remained stable. Importantly, we were comparing a group of individuals aspiring to become professional musicians to a group of amateur musicians who actually have a history of comparable years of playing an instrument but with different intensity and a different goal in mind. This stands in contrast to many other studies that have used nonmusicians as a comparison group. All of our participants look back on similar amounts of musical training, but the aspiring professionals presumably have been trying, for quite some time, to perfect their general ear-training skills in order to pass a highly competitive entrance exam. Accordingly, we found some structural differences between aspiring professionals and amateurs at the beginning of our observation period, with aspiring professionals exhibiting more gray-matter volume in hippocampus, superior parietal lobule, superior/middle temporal gyrus, and postcentral gyrus. However, in the following weeks and months, aspiring professionals actually exhibited a decrease of graymatter volume over time compared to amateur musicians.

At first, the observed decrements in gray-matter volume among aspiring professionals may seem counterintuitive. However, we have argued before that plasticity might in part be characterized by volume expansion followed by a selection process leading to a partial renormalization of overall volume (Wenger, Brozzoli, et al. 2017). In fact, given the large number of skills humans acquire during their lifetime, plasticity cannot be conceived as a process of perpetual growth (Changeux & Dehaene, 1989; Lindenberger et al., 2017; Wenger et al., 2017). According to the exploration–selection–refinement (ESR) model of human brain plasticity (Lindenberger & Lövdén, 2019) neuronal microcircuits potentially capable of implementing the computations needed for executing novel skills are, early in learning, widely probed, with a concomitant increase in gray-matter volume. This phase of exploration is followed by phases of experience-dependent selection and refinement of reinforced microcircuits and the gradual elimination of novel structures associated with unselected circuits. It is tempting to speculate that the aspiring professionals had entered the selection and refinement phases of a plastic episode when they were recruited for participation in the present study. Clearly, this interpretation needs to remain tentative because we did not observe the full cycle of volume expansion followed by renormalization as in our previous study on motor training (Wenger, Kühn, et al. 2017) or as Quallo and colleagues did in their study on tool-use in monkeys (Quallo et al. 2009). Nevertheless, it offers a tenable explanation for the observed structural decreases in left planum polare, posterior insula, and inferior frontal orbital gyrus that needs to be corroborated in future work.

Thus far, data that are consistent with the ESR model have been primarily observed in early ontogeny or during motor skill acquisition; for review, see Lindenberger and Lövdén 2019. Acquiring a complex skill like playing an instrument, in combination with mastering the complexities of harmony and ear training is a different story. There are no data available yet that chart the sequential progression of plasticity over years of musical training. What is documented in the literature are, for the most part, cross-sectional studies showing differences in brain structure between musicians and non-musicians. We can therefore only speculate how the alteration of brain structure in response to years of musical training that has evidently resulted in lasting volume expansion can be reconciled with an ESR view of plastic change. One possibility is that changes occur as a sequence of *several* expansion–renormalization cycles that always conclude in only partial renormalization. This would in the long run result in a building-up of consistently “skill-optimized” gray-matter structure. Obviously, we could not investigate this hypothesis in the current study. What we have observed is a decrease in estimates of gray-matter volume in the group of musicians intensely preparing for an entrance exam, in comparison to a group of musicians still actively performing music on a daily basis but without intensive training. It is noteworthy that others have reported associations between smaller volume and higher expertise: In ballet dancers (Hänggi et al. 2010) and also in skilled pianists (Granert et al. 2011), striatal volume was smaller in individuals with greater motor function efficiency. Furthermore, in a study investigating nonmusicians, amateurs, and expert musicians, there was a negative correlation between degrees of music expertise and graymatter density in right postcentral gyrus, bilateral precuneus/paracentral lobule, left inferior occipital gyrus, and bilateral striatal areas (James et al. 2014).

Following up on our structural results, we also investigated whether we would see indications of plasticity at the functional level. If what we observed here is indeed the second part of an expansion–renormalization cycle, then the left planum polare, which made up the largest patch of gray matter showing volume reduction, would be expected to undergo changes in functional connectivity. Hence we expected that the planum polare would show increased connectivity throughout the brain, specifically to regions previously implicated in musical processing. Indeed, resting-state functional connectivity analyses revealed that over time, the left planum polare was better connected within left auditory cortex itself extending towards superior temporal pole, and also to the right auditory cortex and superior temporal pole, left precentral and also supplementary motor area, left posterior cingulate cortex, and left and right postcentral gyrus, regions that have been shown before to matter in music expertise (Luo et al. 2012; Groussard et al. 2014).

Left auditory cortex has been shown to be involved in processing of melody (Bengtsson and Ullén 2006) and more specifically also in musical semantic memory (Groussard et al. 2010). Left posterior cingulate cortex has been discussed in the context of integrating sensory information and emotional content, for example during reading musical notation (Hyde et al. 2009), in the context of familiarity tasks featuring well-known songs (Satoh et al. 2006), and in combination with autobiographical memories associated with musical excerpts (Ford et al. 2011). Supplementary motor area has been shown before to exhibit greater gray-matter volume in musicians versus nonmusicians (Gaser and Schlaug 2003) and has been implicated in the processing of sequential temporal structures (Bengtsson et al. 2009), pitch and timing repetition during both listening and performance tasks (Brown et al. 2013), as well as in rhythmic and melodic musical improvisation (de Manzano and Ullén 2012).

Also, the results of our graph theoretical analysis go along with our assumption that the region of decreased volume exhibits improved functionality after restructuring and indeed point to the fact that planum polare is now participating in a more complex network. This is reflected in all global measures of graph complexity we investigated, but not on the nodal level.

At all measurement occasions, we observed significant correlations between individual differences in gray-matter volume decrements and music expertise. In other words, the highest performing individuals exhibited the most pronounced decreases in gray-matter volume in left planum polare, left insula, and left inferior frontal gyrus, thus, show the largest plastic change on the neural level. However, counter to expectations, we did not observe any significant correlations between changes in music expertise and changes in gray-matter volume. One reason for the absence of such a change-change association is the high degree of stability of individual differences in music expertise over time. For instance, in aspiring professionals, we observed the following correlations in music expertise between adjacent measurement occasions (*r*_AB_ = .955; *r*_BC_ = .896; *r*_CD_ = .950; *r*_DE_ = .981).

We can only speculate about the neurobiological mechanisms that may have caused the observed reductions in gray-matter volume. Synaptic changes including dendritic branching and axon sprouting as well as glial changes come to mind and we and others have elaborated on the exact potential mechanisms before (Lindenberger & Lövdén, 2019; Wenger, Brozzoli, et al., 2017; Zatorre, Fields, & Johansen-Berg, 2012). Future studies need to incorporate additional MR sequences specifically tailored to disentangle these processes, as for example T_1_ maps (Tardif et al. 2016; Lerch et al. 2017).

The present study also has some further limitations that need be mentioned. First, there was no random assignment of participants to groups. Obviously, this caveat is inherent in the studied topic and is not easy to overcome. We have tried to limit this problem by recruiting two groups of participants with comparable years of playing an instrument. Still, there might be pre-existing differences between people who aspire to become professional musicians and people who consider themselves amateur musicians (Ullén et al. 2016). In addition, the stress to which aspiring professional musicians are exposed might have influenced the present results, as stress has been shown to result in gray-matter volume reductions (Kassem et al. 2013). Thus, we cannot rule out that the observed decreases in gray-matter volume might, to some extent, be related to stress, even though our findings of increased functional connectivity and the correlation with behavioral performance renders this explanation rather unlikely, and also auditory cortex does not belong to those regions typically affected by stress-related reductions (Lupien et al. 2009). Finally, the present samples were not systematically stratified by which main instrument the participants played. Hence, we may have missed out on effects that are specific to particular focal instruments, such as piano versus strings.

To conclude, we found that musicians intensely preparing for the entrance exam to a University of the Arts show reliable reductions in gray-matter volume in regions pertinent to music expertise, whereas a group of amateurs not preparing for an exam did not show such changes. The planum polare, which was the largest gray-matter cluster with volume reductions, showed increasing functional connectivity to other musically-relevant regions. This increase in connectivity was also reflected in global metrics of network participation based on graph theory. The present results are consistent with the ESR model of plastic change (Lindenberger and Lövdén 2019), which posits an expansion of gray-matter volume during early phases of skill acquisition, followed by partial renormalization (Wenger, Brozzoli, et al. 2017).

## Acknowledgements

This work was supported by the Max Planck Institute for Human Development and an intramural grant from the Innovation Fund of the President of the Max Planck Society given to UL. We are grateful for the assistance of the MRI team at the Max Planck Institute for Human Development in Berlin. The authors thank Andreas Brandmaier and Ziyong Lin for setting up the hierarchical structural equation model for the behavioral task data, Caroline Garcia Forlim for advice on the graph theoretical analyses, as well as all participants and student assistants.

## Footnotes

1 Missing data for one participant.

