## Supplemental Figure 1 for "Observing plasticity of the auditory system: Volumetric decreases along with increased functional connectivity in aspiring professional musicians"

### Supplementary material.

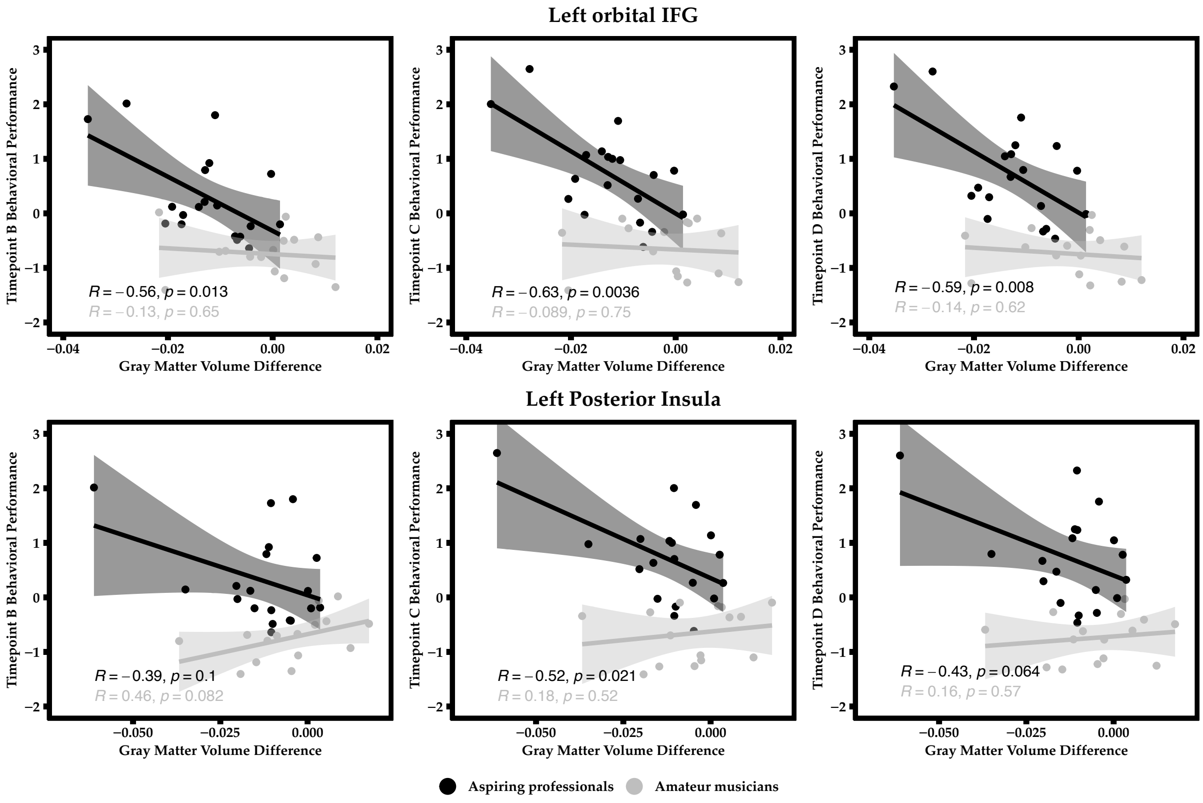


**Figure S1.** Correlation between gray-matter decreases in left orbital inferior frontal gyrus (IFG) and left posterior insula inferior frontal orbital gyrus and behavioural performance at respective measurement occasions B, C, and D.
